# Spatiotemporal ontogeny of brain wiring

**DOI:** 10.1101/385369

**Authors:** Alexandros Goulas, Richard F. Betzel, Claus C. Hilgetag

## Abstract

The wiring of the brain provides the anatomical skeleton for cognition and behavior. Connections among brain regions have a diverse and characteristic strength. This strength heterogeneity is captured by the wiring cost and homophily principles. Moreover, brains have a characteristic global network topology, including modularity and short path lengths. However, the mechanisms underlying the inter-regional wiring principles and global network topology of brains are unknown. Here, we address this issue by modeling the ontogeny of brain connectomes. We demonstrate that spatially embedded and heterochronous neurogenetic gradients, without the need of axonal-guidance molecules or activity-dependent plasticity, can reconstruct the wiring principles and shape the global network topology observed in adult brain connectomes. Thus, two fundamental dimensions, that is, space and time, are key components of a plausible neurodevelopmental mechanism with a universal scope, encompassing vertebrate and invertebrate brains.

## Introduction

Cognition and behavior rely on the communication among distinct parts of the brain, which is mediated by structural connections (1, 2). Empirical studies in vertebrate and invertebrate brains have demonstrated that a graded and highly heterogeneous strength of structural connections is a basic aspect of brain connectomes (3–6). Moreover, brain networks exhibit a characteristic global topology (Fig. 1) (7, 8), including features such as modularity and short path lengths. These network topology features pertain to brains as diverse as the fruit fly and the human brain, and presumably facilitate integration and segregation of the distinct brain regions (7).

**Fig. 1.**
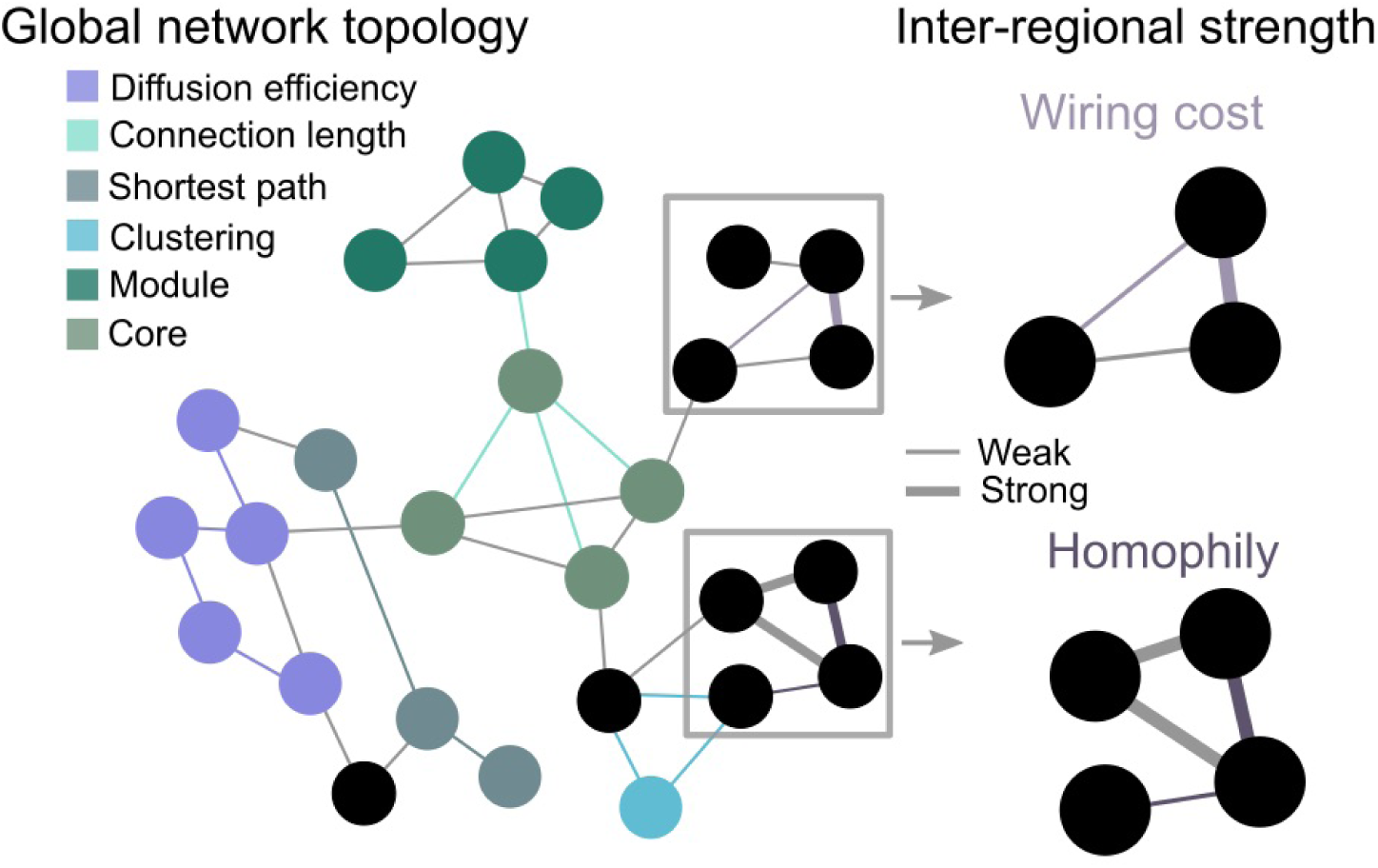
Inter-regional strength heterogeneity (right) and global network topology (left) of brain connectomes. Heterogeneity of inter-regional connectivity strength is parsimoniously described by the wiring cost and homophily principles. Schematic depiction illustrates characteristic global network topology features observed in diverse species. For simplicity, directionality of connections is omitted.

A parsimonious description of the heterogeneous strength of inter-regional connections is offered by the principles of wiring cost and homophily of connections (Fig. 1). These principles pertain to a diversity of species (9–16). The wiring cost principle dictates that brain regions, presumably due to material and metabolic cost constrains, are strongly connected with brain regions that are spatially close, whereas connections with spatially distant brain regions, if any, are very weak (14). Physical distance explains, from a statistical standpoint, a large percentage of connectional strength heterogeneity (14). A more recent principle explaining strength heterogeneity is the homophily of connections. The principle of homophily of connections is based on observations in social networks (17). In social networks, two persons with many common close friends are likely to become close friends themselves. In an analogous fashion, two brain regions with many common brain regions that they strongly connect to, are likely to be themselves strongly connected. (10, 11, 15, 16, 18). Connectional homophily also explains, from a statistical standpoint, a large percentage of connectional strength heterogeneity (11). In sum, empirical evidence demonstrate that the wiring cost principle, as reflected in the physical distance between brain regions, and the homophily principle offer a parsimonious description of the heterogeneous strength of structural connections.

Wiring principles not only offer a parsimonious description of connectomes, but also guide approaches for deciphering the mechanisms underlying these principles. With respect to the homophily principle, tentative mechanistic explanations rely on Hebbian-like plasticity (11, 15, 19). Other views postulate that the connectional homophily of the brain is a consequence of its spatial embedding and the need for functionally similar regions to be interconnected (10). With respect to the wiring cost principle, despite a plethora of studies demonstrating that neural systems are configured in such a way to minimize their wiring cost, the mechanisms that result in such an economical configuration remain elusive. A parsimonious suggested mechanism is based on the spatial embedding of the brain and the mere probabilistic fact that it is more likely for a brain region to establish connections with proximal rather than distant regions (20, 21). With respect to the global network topology of brains, generative models based on empirically deciphered rules can lead to synthetic connectomes with a global network topology that is comparable to the topology of empirical connectomes (16), while other models emphasize the importance of the temporal evolution of networks in shaping certain network properties (22).

In order to gain neurobiologically plausible insights into the ontogeny of brain connectomes, there is a pressing need to move beyond teleological function-based explanations of brain wiring and adopt parsimonious modeling approaches that are interpretable at the neurobiological and mechanistic level. Moreover, there is the need to employ such a modeling approach in a comparative, cross-species context in order to uncover the potentially universal scope of key components of plausible mechanisms. Here, we employ such a modeling approach and directly compare our modeling results with empirical observations in invertebrate and vertebrate brains. First, we use empirical connectomes and offer a comprehensive demonstration of the universality of the connectional homophily and wiring cost principles. Second, we show with computational modeling that spatially ordered and heterochronous neurogenetic gradients, conjointly with stochastic axonal growth, can reconstruct the wiring principles and shape the global network topology observed in empirical connectomes. Our modeling work highlights space and time, that is, spatial embedding and heterochronicity, as key components of a plausible, universal and neurobiologically realistic developmental mechanism for brain wiring.

## Results

### Homophily and wiring cost principles

We analyzed drosophila, mouse, macaque monkey and human connectomes for which quantitative data on strength of connections was available (3–6, 16). The cosine similarity between regions A and B was used as a measure of homophily. Both incoming and outgoing connections were taken into account for the computation of the homophily. For each pair of regions A and B, connections between regions A and B were excluded in computing the homophily of regions A and B. Cosine similarity ranges from 0 to 1, with (0) 1 denoting connectionally (dis)similar regions, thus, exhibiting (low) high homophily. The physical distance of the brain regions was used as a wiring cost metric.

We first used the empirical vertebrate and invertebrate brain connectomes for offering a comprehensive view of the principles of wiring cost and homophily. We binned the homophily and distance values and for each bin we calculated the average strength of connections (Fig. 1). For all connectomes, increased homophily was accompanied by increased connection strength. Distance had the opposite effect, thus, increased distance was accompanied by decreased connection strength (Fig. 2).

**Fig. 2.**
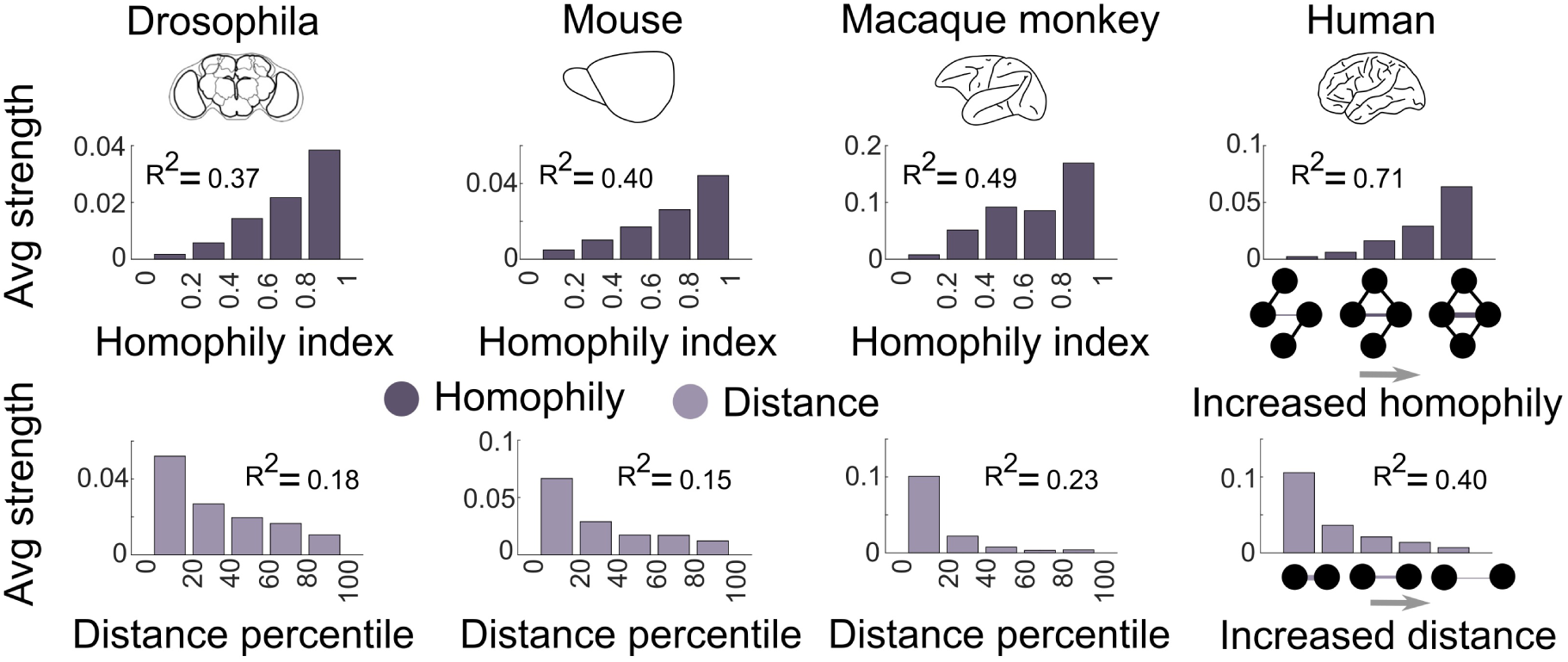
Homophily, physical distance and strength heterogeneity of empirical brain connectomes. Increased homophily entails increase of connectivity strength. Increased physical distance entails decrease of connectivity strength. Depicted *R*^2^ values are derived from univariate regression. See section “Homophily and wiring cost principles” for results and *R*^2^ values concerning multivariate regression.

For a formal statistical assessment of the contribution of homophily and distance to the strength of connections, a least square regression was used (SI Appendix, Materials and Methods). For the drosophila connectome, fitting the homophily and distance to the strength of connections resulted in significant contributions of both factors (univariate model: *β*=0.61,-0.41 p<0.001, model fit *R*^2^=0.37,0.18, multivariate model: *β*=0.53,-0.14 p<0.001, model fit *R*^2^=0.39, for homophily and distance, respectively). Significant results were obtained for the rest of the connectomes, namely for the mouse (univariate model: *β*=0.63,-0.38 p<0.001, *R*^2^=0.40,0.15, multivariate model: *β*=0.58,-0.09 p<0.001, *R*^2^=0.41, for homophily and distance, respectively), macaque monkey (univariate model: *β*=0.69,-0.48 p<0.001, *R*^2^=0.49,0.23, multivariate model: *β*=0.63,-0.12 p<0.001, *R*^2^=0.50, for homophily and distance, respectively) and human (univariate model: *β*=0.84,-0.62, p<0.001, *R*^2^=0.71,0.40, multivariate model: *β*=0.79,-0.10, p<0.001, *R*^2^=0.71, for homophily and distance, respectively). These aforementioned empirical model fit values were used as a measure for the quality of the fit of the synthetic connectomes to the empirical connectomes (see section “Comparing wiring principles in synthetic and empirical connectomes”).

In sum, in both vertebrate and invertebrate brains, homophily and wiring cost constitute wiring principles that capture a substantial part of the heterogeneity of inter-regional connectivity strength. Increasing physical distance dictates decreased strength of connections and increasing homophily dictates increased strength of connections.

### Simulation of neurogenetic gradients and connectivity formation

Heterochronicity in development was suggested to play an important role in the morphology of animals (23). In addition, heterochronous neurogenesis seems related to the preferential connectional patterns between brain regions of the rat and mouse brain (12, 24). Inspired by such theoretical work and empirical evidence, we aimed at simulating neurogenesis and connectivity formation in a model based on heterochronicity (see SI Appendix, Materials and Methods). A 2D synthetic brain constituted the niche of the simulations, with each unit surface hosting up to *N* neurons (Fig. 3 A). Heterochronous neurogenesis emanated from neurogenetic root(s)/origin(s), forming orderly, spatially embedded, neurogenetic gradients, as empirical evidence dictate (25, 26) (Fig. 3 A). Each unit surface of the synthetic brain was assigned a time window dictating the probabilities of a neuronal population of *n* neurons migrating at each unit surface at a given time point *t* of the simulation (Fig. 3 A). This time window was proportional to the physical distance of each unit surface from the neurogenetic root(s). This modeling choice was informed by empirical evidence suggesting spatially ordered waves of migration of neurons from neurogenetic origins (25, 26). When a neuron migrated to a unit surface, it attempted to form a connection by extending an axon towards a direction randomly sampled from a uniform distribution. If the straight line of this axon extension intersected a circle with a radius *r* centered around each neuron that have already been placed to the synthetic brain, then a connection was formed (see SI Appendix, Materials and Methods) (Fig. 3 B). A straight line was used, as in previous modeling approaches (20, 27), in the absence of an empirical model capturing the exact trajectory of developing axons in vertebrate and invertebrate brains.

**Fig. 3.**
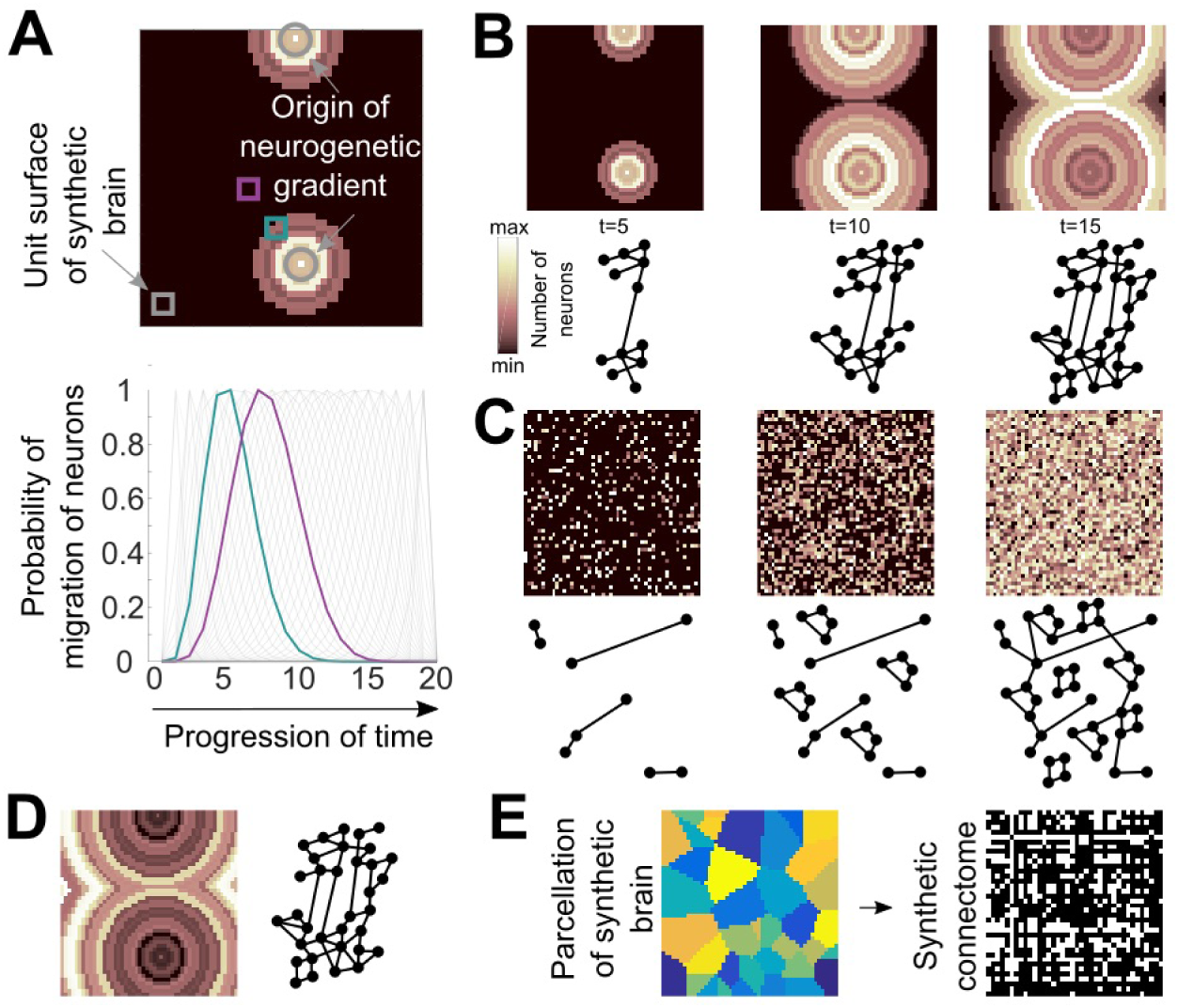
Developmental modeling approach. A. Neurogenetic gradients were simulated in a synthetic 2D brain. Each surface unit was characterized by a time window that indicates at a given time point the probability of a neuronal population migrating to each surface unit. Each time window is shaped by the distance of each surface unit from the root(s), that is, origin(s), of the neurogenetic gradients. For instance, the surface unit close to the neurogenetic origin (petrol green) is more probable to be populated earlier than the surface unit further from the neurogenetic origin (magenta). B. Heterochronous and spatially ordered ontogeny of synthetic connectomes. C. Heterochronous, spatially random and D. tautochronous ontogeny of synthetic connectomes. E. Creation of the synthetic connectome.

For assessing the importance of heterochronicity and the spatial order of neurogenesis, two additional scenarios were simulated. In one scenario, heterochronous neurogenesis and connectivity formation was simulated as previously described, but the spatial manifestation of neurogenesis was random (Fig. 3 C). In the other scenario, neurogenesis was tautochronous, that is, all neurons were simultaneously placed in the synthetic brain and formed connections (Fig. 3 D). Importantly, in all scenarios, the final synthetic brain contained the exact number of neurons, thus, potential differences in model fit could not be attributed to this factor.

Since the empirical connectomes are region-to-region connectivity matrices, the synthetic brain was parcellated into regions with a Voronoi tessellation resulting in a synthetic region-to-region connectivity matrix (Fig. 3 E). The number of regions and connectivity density, that is, the fraction of connections that exist over the total number of connections that can exist given a number of regions, of the synthetic connectomes was determined based on the number of regions and connectivity density of the empirical connectomes. Connection strength was calculated as the number of projections from region A to B divided by A’s total number of projections. This normalization renders the connectomes comparable across simulations, empirical data, as well as across species.

### Comparing wiring principles in synthetic and empirical connectomes

The synthetic connectomes obtained from the simulations corresponding to each scenario (heterochronous spatially ordered, heterochronous spatially random, tautochronous), were subject to a least squares regression analysis in order to obtain the parameters (*β* values) that correspond to the statistical association of homophily and distance to the strength of connections. These parameters, obtained exclusively from the synthetic connectomes, were subsequently used to estimate the fit to the empirical connectomes by predicting the strength of the connections of the empirical connectomes.

### Model selection

Akaike’s information criterion (AIC), controlling for the complexity and fit of models, was used for ranking the models. We found that the heterochronous better relate to the homophily principle and tautochronous models better to the distance-based wiring principle, when these principles were examined separately. Importantly, the heterochronous and spatially ordered model was better when considering the two principles simultaneously. Specifically, for the homophily principle, the heterochronous models resulted in the lowest AIC values for all species (Figs. S1 and S2). The temporal overlap of neurogenesis, controlled by parameter *a*, and the number of origins of the neurogenetic gradients did not systematically influence the AIC values (Fig. S1), thus, average summaries for each scenario are provided (Fig. S2). For the wiring cost principle, the tautochronous model resulted in the lowest AIC values for all species (Figs. S1 and S2). Importantly, when considering simultaneously the homophily and wiring cost principles, since both co-characterize adult brain connectomes, the heterochronous and spatially ordered model exhibited the lowest AIC values (Figs. S1 and S2).

### Model fit and null models

AIC is useful for ranking candidate models, but it does not offer information on the quality of the fit of the models and how different, from a statistical standpoint, such fit is in relation to a null model. Therefore, we used the coefficient of determination (*R*^2^) that is based on the residual sum of squares of connection strength, as predicted from the parameters obtained exclusively from the synthetic connectomes, and the actual, empirical connection strength. An example of such empirical and predicted strength of connections is provided in Fig. S3. Importantly, the *R*^2^ values obtained from the fit of the parameters from the synthetic connectomes to the empirical connectomes were compared with the *R*^2^ values obtained from the parameters and fit when considering only the empirical connectomes. Moreover, the *R*^2^ offered the possibility of statistical null hypothesis testing (*F*-test).

With respect to the homophily principle, the best fit to the empirical connectomes for all species was observed for the heterochronous models (median *R*^2^=0.37, *R*^2^=0.39, *R*^2^=0.48 and *R*^2^=0.68, for the drosophila, mouse, macaque monkey and human connectomes, respectively) (Fig. 4). The parameter denoting the importance of homophily in predicting the strength of connections obtained from the synthetic models resulted in predictions of empirical connectivity with a fit that was very close to the fit obtained from a model fitted exclusively to the empirical connectomes (*R*^2^=0.37, *R*^2^=0.40, *R*^2^=0.49 and *R*^2^=0.71, for the drosophila, mouse, macaque monkey and human connectomes, respectively) (Fig. 4). Hence, heterochronously developed and wired brains resulted in 98% (median) of the empirical fit (range 96-100%) that corresponds to the relation between connectivity and homophily in empirical connectomes (Fig. 4).

**Fig. 4.**
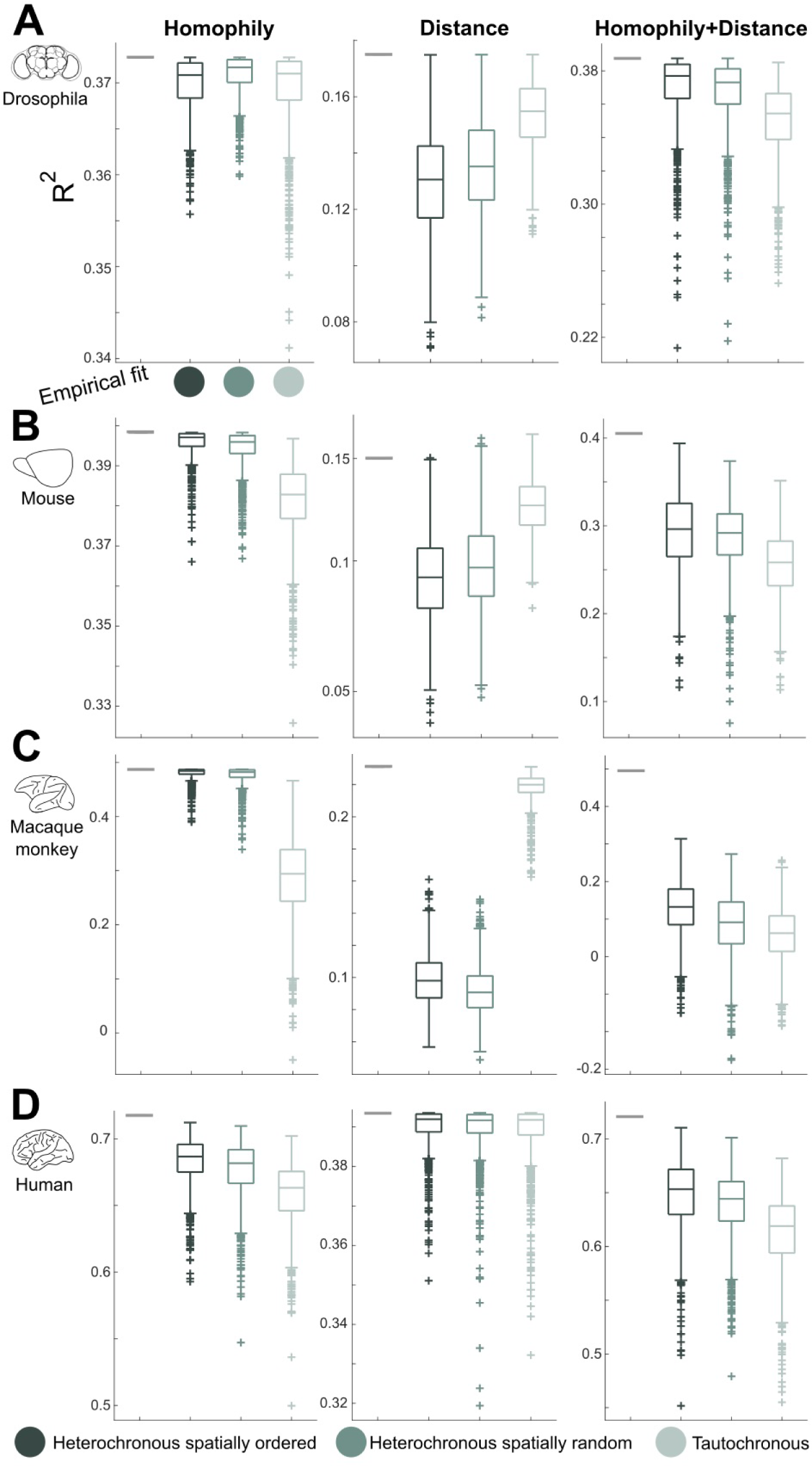
Inter-regional wiring principles captured by the synthetic connectomes. Summary of *R*^2^ values for the predictions of empirical connectivity from parameters calculated exclusively from the synthetic connectomes. Values for *R*^2^ are depicted for the A. drosophila, B. mouse, C. macaque monkey and D. human connectomes when considering the homophily or wiring cost principles separately or simultaneously. Note the better fit of the heterochronously generated connectomes when considering simultaneously the wiring cost and homophily principles. Note that the predictions from model parameters estimated exclusively from the synthetic connectomes, result in *R*^2^ values very close to *R*^2^ values obtained when using exclusively the empirical data (grey bars), thus, replicating a high percentage of the empirically observed relations between connectivity strength, homophily and distance across vertebrate and invertebrate brains.

With respect to the wiring cost principle, the best fit to the empirical connectomes for all species was observed for the tautochronous models (median *R*^2^=0.15, *R*^2^=0.12, *R*^2^=0.22, *R*^2^=0.39, for the drosophila, mouse, macaque monkey and human connectomes, respectively) (Fig. 4). The parameter denoting the importance of distance in predicting the strength of connections obtained from the synthetic models resulted in a fit to the empirical connectomes that was very close to the fit obtained from a model fitted exclusively to the empirical connectomes (*R*^2^=0.18, *R*^2^=0.15, *R*^2^=0.23, *R*^2^=0.40, for the drosophila, mouse, macaque monkey and human connectomes, respectively) (Fig. 4). Hence, tautochronously developed brains resulted in 89% (median) of the empirical fit (range 80-98%) that corresponds to the empirical relation between connectivity and distance (Fig. 4).

Importantly, when considering the homophily and wiring cost principles simultaneously, since both co-characterize empirical connectomes, the heterochronous and spatially ordered model resulted in the best fit across all species (*R*^2^=0.37, *R*^2^=0.29, *R*^2^=0.13, *R*^2^=0.65, for the drosophila, mouse, macaque monkey and human connectomes, respectively) (Fig. 4). Synthetic brains that were developed in an heterochronous and spatially ordered manner resulted in parameters, denoting the importance of wiring cost and homophily principles in predicting the strength of connections, that led to a fit to the empirical connectomes that was very close to the fit obtained from a model applied exclusively to the empirical connectomes (*R*^2^=0.39, *R*^2^=0.41, *R*^2^=0.50, *R*^2^=0.71 for the drosophila, mouse, macaque monkey and human connectomes, respectively) (Fig. 4). Therefore, for the simultaneous assessment of the homophily and wiring cost principles, brains that were developed and wired in an heterochronous and spatially ordered fashion resulted in almost the same fit obtained directly from the empirical connectomes, reaching 81% (median) of the empirical fit (range 26-95%) (Fig. 4). Lastly, all the predictions of the strength of the empirical connections from the parameters obtained from the synthetic connectomes were statistically significant (p<0.001, *F*-test) (Table S1).

In all, despite the fact that the simulations were completely agnostic to the principles that characterize empirical connectomes, the synthetic connectomes replicated a high percentage of the empirical relation between homophily, distance and connectivity strength, with the heterochronous and spatially ordered model exhibiting superior fit when considering both wiring principles simultaneously. Explaining the residual part of the empirical fit not captured by our model (Fig. 4) may involve additional mechanisms, such as pruning, axonal guidance or activity-dependent plasticity.

### Ontogeny of global network topology

After the examination of inter-regional wiring principles, we next sought to investigate to what extent the synthetic connectomes generated under the different developmental scenarios reflected differences in global network topology. We focused on widely and commonly used global network topology metrics, that is, clustering coefficient, characteristic path length, diffusion efficiency, physical distance spanned by the connections, maximum clique size (defining the core of the network), and modularity (7, 8, 14, 28).

First, for each species separately, we constructed a network topology profile for each of the synthetic connectomes that were generated under the three different scenarios, the later defining three different groups. We performed a canonical discriminant analysis (CDA) to investigate if the synthetic connectomes were distinct in terms of their network topology profile and what network topology metrics are the most prominent arbiters for such a distinction. The components obtained from the CDA defined a network topology morphospace (29) that separated the synthetic connectomes generated from the three different scenarios for all species (Wilks’ Λ=0.24, 0.21, 0.13, 0.43, all p<0.001, permutation test, for the drosophila, mouse, macaque monkey and human connectomes, respectively) (Fig. 5 A). Nearly all between-groups variance was explained by the first component (82%, 95%, 98%, and 98%, for the drosophila, mouse, macaque monkey and human connectomes, respectively), that separated the heterochronously from the tautochronously generated synthetic connectomes (Fig. 5 A). Overall, heterochronously generated connectomes exhibited higher network topology metrics that differentiated them from the tautochronously generated connectomes, with the size of the core (max clique size) and characteristic path length constituting the most prominent network metrics segregating the synthetic connectomes in the network topology morphospace (Fig. 5 A).

**Fig. 5.**
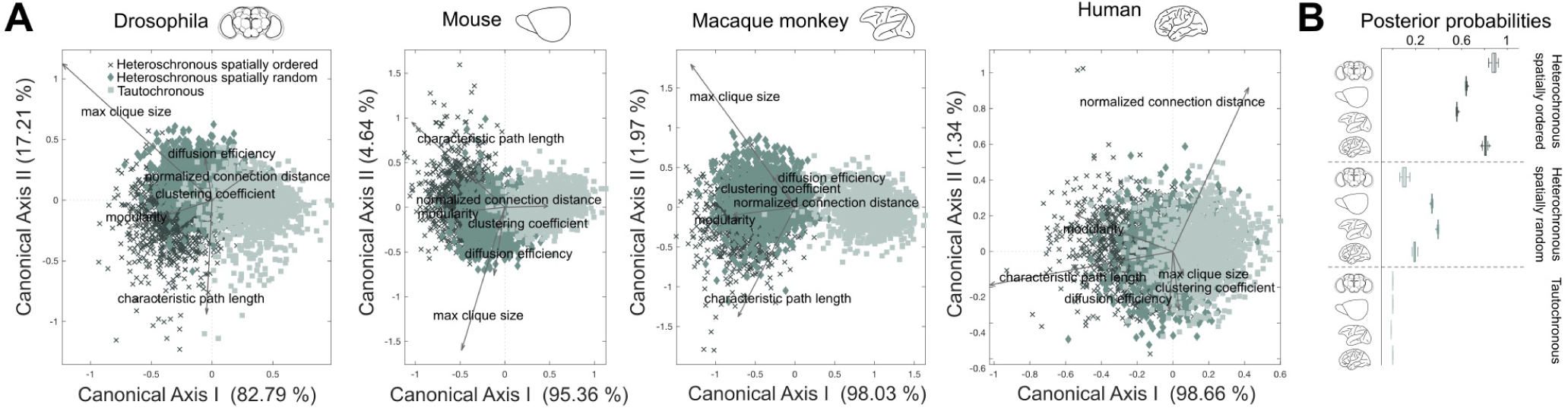
Network topology morphospace of synthetic connectomes and classification of empirical connectomes. A. Heterochronously and tautochronously generated connectomes are separated based on their global network properties across the first component. Core size (max clique size) and characteristic path length are the major network metrics differentiating the groups. B. Posterior probabilities indicate that the global network topology signature of all empirical connectomes are more reminiscent of connectomes with an heterochronous ontogeny.

Next, we examined if the global network topology of the empirical connectomes mostly resembled the topology of heterochronously or tautochronously generated synthetic connectomes. To this end, we situated the empirical connectomes in the network topology morphospace defined by the heterochronously and tautochronously generated connectomes. This was achieved by performing the CDA 100 times, each time with 70% of the synthetic connectomes, and using the derived model parameters to calculate the posterior probabilities of the empirical connectome belonging to one of the three groups. This procedure was repeated for each species separately. The higher posterior probabilities were obtained for the connectomes that were developed in an heterochronous and spatially ordered way (Fig. 5 B).

Empirical connectomes mostly resemble the heterochronous and spatially ordered synthetic connectomes and are maximally dissimilar to random connectomes (Fig. S4). However, empirical connectomes are also distinct from the synthetic connectomes (Fig. S4), indicating that certain empirical network metrics are not accurately captured by the synthetic connectomes. We assessed how faithfully the synthetic connectomes reconstructed the empirical global network topology by plotting each network metric separately for the synthetic and empirical connectomes. Overall, good correspondence with empirical values was observed, with certain deviations, either underestimating or overestimating the empirically observed values of the network metrics (median % deviations from 100% corresponding to a perfect reconstruction): 9%, 34%, 32%, 13%, for the drosophila, mouse, macaque monkey and human connectomes, respectively) (Fig. S5). Therefore, although that the global network topology of the empirical connectomes is more reminiscent of connectomes that were developed in an heterochronous and spatially ordered fashion, deviations indicate that a more accurate reconstruction of certain empirical global network topology aspects may involve additional mechanisms, such as pruning, axonal guidance or activity-dependent plasticity.

### Discussion

Heterochronicity in development has been suggested as a key underlying principle of the observed morphological diversity in animals (23) and seems to underlie the intricate connectional configuration of rodent brains (12, 24). Here, we brought the concept of heterochronicity to the connectional realm, within a comparative framework that encompasses a diverse spectrum of brains, ranging from fruit fly to human brains. We have demonstrated that heterochronous and spatially embedded development and connectivity formation constitutes a key component of a universal and neurobiologically realistic mechanism that may underly the empirically observed inter-regional wiring principles and global network topology of adult brains.

Our empirical results demonstrate that the homophily and wiring cost principle characterizes the adult connectional configuration not only of the mammalian brain (10, 11, 15), but also the insectivore brain. Importantly, our modeling work demonstrates that such a wiring configuration can be laid down by the presence of spatially ordered and heterochronous neurodevelopmental events and a purely stochastic axonal growth, without gradients of axonal guidance molecules or activity-dependent remodeling of connections. Therefore, our results offer a solution to the conundrum of how two regions with high homophily, that is, similar patterns of connections, and thus, functional similarity, end up preferentially and strongly interconnected in the adult brain. Heterochronous and spatially ordered development renders feasible the emergence of such wiring configuration, since two regions that are populated by neurons at proximal time points in development, have similar available connectional targets with other regions, also occupying similar time points in development, leading to high homophily of the two regions, and to the presence of strong connections in-between the two regions. In other words, we show how a structural skeleton that allows connectionally and functionally similar regions to be interconnected can be laid down with a parsimonious neurodevelopmental mechanism. Such connectional configuration may be further refined through subsequent experience-dependent plasticity. Therefore, the current modeling work complements prior modeling work that aims at offering a compact representation of brain wiring, rather than highlighting key components of plausible mechanisms pertaining to the ontogeny of connectomes (30).

Our modeling work also offers a parsimonious mechanism for the well-established wiring cost principle that pertains to diverse connectomes (7). The spatial embedding of the brain and the mere probabilistic fact that the establishment of connections is more likely to take place with proximal rather than distant targets (21), is sufficient to replicate a large part of the empirically observed relation between physical distance and strength of connections, from the drosophila to the human connectome. Thus, our results extend previous modeling and theoretical work (20, 21) by demonstrating a basic, plausible and universal mechanism leading to economically wired brains.

Brain connectomes exhibit non-random global topological properties, such as modularity and short path lengths (7). Connectomes that have an heterochronous ontogeny exhibit a global network topology different from connectomes with tautochronous ontogeny and are distinguishable from each other in a global network morphospace. Importantly, the global network configuration of the brain of diverse species, from drosophila to humans, is mostly reminiscent of connectomes with an heterochronous and spatially ordered ontogeny. Thus, heterochronicity may shape the empirically observed global network topology signature of brains. Deviant connectome topologies are observed in a number of pathologies (31). At the molecular level, the *Notch* and *Bone Morphogenetic Protein* signaling pathways are involved in the initiation of neurogenesis (32). Our results show that pathologies involving these pathways, and consequently the normative temporal unfolding of neurogenesis across the brain, might also affect the global large-scale connectional configuration of the brain.

Our model adopts a parsimonious approach to connectivity formation and replicated a large percentage of empirically observed inter-regional wiring principles and global network topologies. However, these results do not entail that other aspects, such as principles of chemotaxis, that is, chemoattraction and chemorepulsion for guiding axons to their proper targets (33), are irrelevant. Such principles can be incorporated in future extensions of the model in order to reconstruct the currently residual part of the empirically observed inter-regional wiring principles and global network topologies of brain connectomes.

In all, we have demonstrated that heterochronicity is a plausible key universal developmental phenomenon that can sculpt the intricate connectional configuration of vertebrate and invertebrate brains.

## Materials and Methods

Connectivity datasets are freely available from the references cited in the text (see SI Appendix, Supplementary Materials, Connectivity Datasets). For simulation of neurogenesis, connectivity formation, and statistical analysis see SI Appendix, Supplementary Materials, Developmental Modeling and Statistical Analysis and Model Fitting sections.

## Acknowledgments

The authors would like to thank Petra Vértes for helpful comments. Support by a Humboldt Research Fellowship from the Alexander von Humboldt Foundation (A.G.) as well as grants from DGF SFB 936/A1, Z3, and TRR 169/A2 (C.C.H.) is gratefully acknowledged.

